# Alternative splicing preferentially increases transcript diversity associated with stress responses in the extremophyte *Schrenkiella parvula*

**DOI:** 10.1101/2022.10.13.512046

**Authors:** Chathura Wijesinghege, Kieu-Nga Tran, Maheshi Dassanayake

## Abstract

Alternative splicing extends the coding potential of genomes by creating multiple isoforms from one gene. Isoforms can render transcript specificity and diversity to initiate multiple responses required during transcriptome adjustments in stressed environments. Although the prevalence of alternative splicing is widely recognized, how diverse isoforms facilitate stress adaptation in plants that thrive in extreme environments are unexplored. Here we examine how an extremophyte model, *Schrenkiella parvula*, coordinates alternative splicing in response to high salinity compared to a salt-stress sensitive model, *Arabidopsis thaliana*. We use Iso-Seq to generate full length reference transcripts and RNA-seq to quantify differential isoform usage in response to salinity changes. We find that single-copy orthologs where *S. parvula* has a higher number of isoforms than A. thaliana as well as S. parvula genes observed and predicted using machine learning to have multiple isoforms are enriched in stress associated functions. Genes that showed differential isoform usage were largely mutually exclusive from genes that were differentially expressed in response to salt. *S. parvula* transcriptomes maintained specificity in isoform usage assessed via a measure of expression disorderdness during transcriptome reprogramming under salt. Our study adds a novel resource and insight to study plant stress tolerance evolved in extreme environments.

## Introduction

Alternative splicing produces different mature RNAs from a single gene. Its impact on increasing transcript diversity has continued to broaden our understanding of gene regulatory mechanisms since it was first observed in 1977 ^1,2^. The potential to create novel transcript diversity via alternative splicing is immense. The Drosophila *DSCAM* gene, which functions as an axon guidance receptor, is an extreme example of alternative splicing. It contains 115 exons and is estimated to give rise to more than 38,000 isoforms that are spatiotemporally regulated to achieve specific regulation in Drosophila neural development ^3,4^. Alternative splicing events can be observed in more than 95% of human genes ^5^. High throughput proteomics and ribosome bound mRNA sequencing (Ribo-Seq) studies show that a significant fraction of alternative splice variants are translated into protein isoforms ^6,7^. Additionally, an ever-increasing array of transcriptome sequencing has revealed the existence of novel non-coding RNAs generated through alternative splicing suggesting their importance in gene regulatory circuits in all eukaryotic clades ^8,9^. Differential splicing in closely related species have shown to reflect their divergent adaptive strategies not readily detectable at the primary gene expression level ^10^. Tissue- and species-specific splicing is more divergent than promoter level divergence among closely related species facilitating independent evolutionary trajectories fitting to each species as highlighted by Calarco et al.^11^ Therefore, genome wide discovery of new transcripts produced via alternative splicing becomes a critical initiative to understand the diversity of gene products and systematically assess their role in evolutionary innovations.

Alternative splicing increases proteome diversity as well as regulatory complexity in plants ^12–14^. In the model plant *Arabidopsis thaliana,* majority of the genes (>60%) undergo alternative splicing. There are more than 70,000 non-redundant transcript isoforms reported for *A. thaliana* ^15,16^. Similar reports on maize ^17^, sorghum ^18^, and cotton ^19^ demonstrate that alternative splicing is prevalent in plants. Differential splicing has also been targeted in crop breeding as shown with sunflowers ^20^.

Large scale changes in alternative splicing have been reported to allow transcriptional adjustments in response to abiotic stresses including salt ^21^, cold ^22^, hypoxia ^23^, and heat stress ^24^. Targeted functional studies have also highlighted the significance of alternative splicing in responses to abiotic stresses. For example, the heat shock protein gene, *hsf2* in *A. thaliana* produces an alternatively spliced transcript resulting in a truncated protein that in turns binds to the *hsf2* promoter to enhance transcription of *hsf2* during heat stress ^25^. While the majority of published studies converge on alternative splicing being a key mechanism for environmental stress adaptation in plants, such studies are limited to abiotic stress sensitive crop models or to *A. thaliana*.

Compared to crop plants, extremophytes that are naturally found in extreme environments are equipped with evolutionary innovations that give them the ability to cope with multiple and extreme levels of environmental stresses ^26^. Therefore, extremophytes could show how the expanded transcriptome diversity via alternative splicing may render additional paths for abiotic stress adaptations absent in stress sensitive models. In this study, we have used the extremophyte model, *Schrenkiella parvula* (formerly *Thellungiella parvula* and *Eutrema parvulum*)^27,28^. to examine its transcriptome diversity augmented by alternative splice variants. *S. parvula* shares a highly co-linear genome with *A. thaliana* ^29,30^. Yet, *S. parvula* is uniquely adapted to multiple abiotic stresses reflecting its natural habitats often associated with hypersaline lakes in the Irano-Turanian region ^31,32^.

Previous studies have shown that the *S. parvula* genome is enriched with duplicated genes associated with abiotic stress responses and stress responsive genes show constitutive high expression as a stress preadaptation compared to *A. thaliana* ^29,32,33^. Alternative splicing plays a complementary role to gene duplications and provides an additional path to increase transcript diversity ^34^. Therefore, we aimed to test the overall hypothesis that alternative splicing leads to increased diversity of stress responsive transcripts in *S. parvula*.

In this study we investigated the complexity of the alternative splicing landscape in roots and shoots in response to salt stress and how alternative splicing may provide transcript diversity associated with adaptations to environmental stress in the model extremophyte *S. parvula.* We used PacBio Iso-Seq sequencing to identify and annotate alternative splice variants and Illumina short reads to quantify isoform abundance. We find that the *S. parvula* transcriptome is enriched in stress-associated isoforms. It shows specific isoform usage in a less disordered state compared to the stress-sensitive model *A. thaliana* in response to high salinity.

## Materials and Methods

### Plant material

*Schrenkiella parvula* (ecotype Lake Tuz, Turkey; Arabidopsis Biological Resource 575 Center/ABRC germplasm CS22663) seeds were grown hydroponically as previously described ^33^. Briefly, plants were grown at a light/dark cycle of 12/12 hr, 100 - 120 mM·m^-2^s^-1^ photon intensity, 20-22 °C, and 1/5^th^ Hoagland’s solution for four weeks. These were treated with a with a combination of 250 mM NaCl, 250 mM KCl, 30 mM LiCl, and 15 mM H_3_BO_3_ for three days to generate tissue samples used to create a reference transcriptome with PacBio Iso-seq sequencing. Shoots and roots were harvested separately. RNA was extracted using QIAGEN RNeasy Plant Mini Kit (QIAGEN, Hilden, Germany) with column digestion to remove DNA contamination. About 4 μg of total RNA per tissue type at a quality of RNA integrity number ≥ 8 based on a Agilent 2100 Bioanalyzer (Agilent Technologies, CA, USA) were used to generate RNA-Seq libraries.

Shoot and root (1 μg) extracted as described above were used for cDNA synthesis using the SuperScript cDNA Synthesis Kit (Invitrogen, Massachusetts, USA) following manufacturer’s instructions to test the presence of multiple isoforms independent from Iso-Seq for a randomly selected gene set expected to express multiple isoforms. Isoform specific PCR primers (Supplementary Table 1) that span the alternative splice sites were designed to use with an amplification protocol (initial denaturation at 95 °C for 3 min; 30 cycles of 95 °C for 30 s, 50-56 °C for 30 s, 72 °C for 2.30 to 3 min; 72 °C for 10 min) run on a Bio-Rad T100 Thermal Cycler (Hercules, CA, USA) with a PCR Master mix Solution i-MAX II (iNtRON Biotechnology, S Korea). PCR products were separated on a 1% agarose gel.

To quantify isoform abundance in response to high salinity compared to control conditions, RNA was extracted from hydroponically grown *S. parvula* and *A. thaliana* (Col-0) as described in Tran et al. (2021). These plants were treated with 150 mM NaCl for 24 hours and harvested together with samples hydroponically grown without added NaCl as a control condition. The hydroponic growth conditions except for the specific salt treatment was kept equivalent to growth conditions given to plants used for reference transcript generation with Iso-Seq. At least 5 plants were used per biological replicate and three biological replicates were used for each root and shoot sample for *S. parvula* and *A. thaliana* to yield a minimum of 1 μg of total RNA per sample used for standard RNA-seq library preparation.

### Transcriptome sequencing

For Iso-Seq based long read sequencing, cDNA synthesis, sequencing library preparation, and PacBio sequencing were conducted at the Arizona Genomics Institute, University of Arizona, USA. Two Iso-seq sequencing SMRT libraries were constructed following size selection from ≤ 4 kb and ≥ 3.5 kb per each tissue and ran on two Pacific Biosciences Sequel cells with v2.1 Chemistry. For RNA-seq based short read sequencing, mRNA enriched cDNA synthesis, library preparation, and sequencing were conducted at the Roy K. Carver Biotechnology Center, University of Illinois Urbana-Champaign, USA. Briefly, True-Seq strand specific libraries (Illumina, San Diego, CA, USA) were multiplexed and sequenced on an Illumina HiSeq4000 platform to generate >15 million 50-nucleotide single-end reads per sample.

### Identification and annotation of full-length transcript models for *S. parvula*

Raw Sequel data were processed using isoseq_sa5.1 pipeline (https://github.com/PacificBiosciences/IsoSeq_SA3nUP. Circular consensus sequences (CCS) were generated from subread BAM files with following parameters: minLength□=□50, -noPolish --minLength=50, --maxLength=15000, --minPasses=1, --minPredictedAccuracy=0.8, --minZScore=-999 --maxDropFraction=0.8. CCS reads were selected as full length reads if it contained the 5’ and 3’ primers and a poly(A) signal preceding the 3’ primer without additional copies of adapters. The full length consensus transcripts were further clustered using ICE (Iterative Clustering for Error Correction) to obtain high-quality isoforms with post-correction accuracy above 99% using Quiver. Error corrected full length reads were mapped to the *Schrenkiella parvula* reference genome v2.2 (Phytozome genome ID: 574) to annotate isoforms assigned to gene models and further select a set of high confidence transcript models. An isoform is annotated as a full length transcript mapped to a genomic locus that has a single gene model assigned. If more than one isoform is mapped to a gene model, the second and subsequent isoforms are considered products of alternative splicing. First, TAPIS ^17^ was used to map isoforms and further error correct the isoforms. To map reads to the genome GMAP ^36^ was used with parameters, --no-chimeras, --cross-species --expand-offsets 1, -K 3000. Then, SQANTI ^37^ was used with default parameters to identify the isoform that matched the primary gene model in the genome and to assign additional isoforms that may be derived from that gene model as alternatively spliced isoforms if both types of full-length isoforms were present in our processed full length data. Canonical splice sites were defined as AG at the acceptor site and GT at the donor site. All the other splice sites were categorized as non-canonical splice sites. Custom python script was used to count canonical and non-canonical splice sites. Finally, we selected non-redundant structurally distinct isoform models that also contained a complete and uninterrupted open reading frame as a selected set of putative protein coding transcript models. Only isoforms that are likely to code for proteins were used for downstream analyses in the current study due to the high uncertainty of functional significance and limited annotation resources available for newly identified non-coding isoforms.

Functional annotations were assigned using PANTHER ^38^ and *A. thaliana* Gene Ontology (GO) annotations (version release date 2020-07-16; DOI:10.5281/zenodo.3954044). Test for enriched functions were performed using BiNGO ^39^. Further clustering of enriched functions were performed using GOMCL ^40^ (with parameters: -gosize 1500 -Ct 0.7 -I 1.5 -hm -nw -d -hg 0 -hgt –ssd) to get a non-redundant set of representative functional annotations at p-values ≤0.05 adjusted for false discovery rate.

### Transcript and gene expression quantification

Following quality checks using FastQC (http://www.bioinformatics.babraham.ac.uk/projects/fastqc/). RNA-seq reads were mapped to gene models for *A. thaliana* (TAIR10) or *S. parvula* (Reference v2.2) as well as transcript models obtained from AtRTD2 ^41^ or *S. parvula* Iso-seq supplemented transcript models using Salmon ^42^ with parameters, “--type quasi -k 31” for indexing and“--gcBias -l A” for quantification. Ortholog pairs between *S. parvula* and *A. thaliana* were assigned based on Oh & Dassanayake 2019 ^43^. RNA-seq reads mapped to gene models were used to identify differently expressed genes. A custom python script was used to count uniquely mapped reads to each gene model. Differentially expressed genes between control and salt treatments within each species were identified using DESeq2 ^44^ RNA-seq reads mapped to transcript models were used for generating expression values for isoforms as well as quantify alternative splicing event frequency. Expression counts for isoforms were converted to TPM (Transcript Per Million) and in comparisons where an isoform was counted as expressed had ≥ 0.5 TPM normalized expression per isoform independent from the expression quantified at the gene level. Isoform ratio per ortholog pairs was calculated based on the number of isoforms per *S. parvula* ortholog divided the number of isoforms detected in the *A. thaliana* ortholog.

Differential splicing was assessed using SUPPA2 ^45^. Briefly, alternative splice events were identified using generateEvents program and differential isoform expression was calculated based on the total expressed number of isoforms per gene using psiPerIsoform included in SUPPA2 together with diffSplice to compare differences in isoform expression between two conditions.

### Shannon entropy calculation for isoform specific transcriptome responses

Isoform expression shifts between conditions or species were quantified using PSI values (proportion of spliced isoforms) assigned for each alternatively spliced isoform per gene as given in the equation below. We used the PSI values to calculate Shannon entropy per gene as described by Ritchie et al. (2008)^46^ and used normalized values between 0 and 1 for between species comparisons as described in Kumar et al. (1986) ^47^

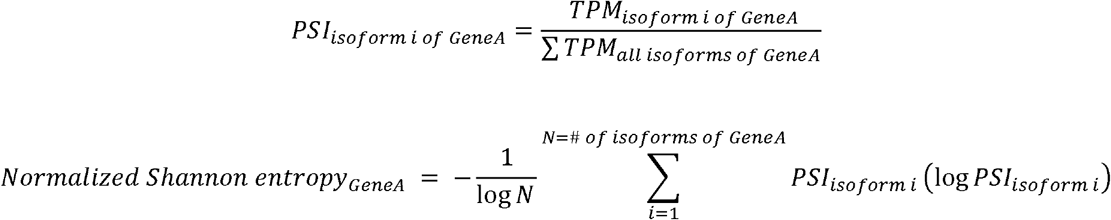

We calculated Shannon entropy values for genes expressed in control and salt treated samples for *S. parvula* and *A. thaliana.* Genes with PSI values less than 0.01 or higher than 0.99 (expected when an isoform is rarely expressed or dominates approximating zero alternative splicing for that gene) were removed from our analysis to test for isoform expression shifts. Further, gene models which were not represented by at least two isoforms were removed from the analysis.

### Splice site prediction for the *S. parvula* genome

We used a deep-neural network, SpliceAi ^48^ to predict genome wide splice sites for *S. parvula* from primary gene model sequences. The network model was trained first with *A. thaliana* gene models from chromosome 1 to 4 and validated with chromosome 5 gene models described in Araport11 ^49^. We provided 200 nucleotides upstream and downstream of a given base scanning all bases per gene in all genes models to predict whether that site is a splice site donor, acceptor or not a splice site. Model prediction was assigned a probability score between 0 and 1 for a given site with values closer to 1 representing the probability of that site being a splice site. We used a probability score of ≥0.6 for the selection of potential splice sites. We used this trained network to predict splice sites for the *S. parvula* genome v2. We compared the predicted splice sites to observed splice sites and identified new splice sites. If new splice sites were predicted for a gene model we had identified more than one isoform, the prediction of a novel splice site or sites for that gene was considered as one additional predicted putative isoform.

## Results

### Improvement of isoform annotation in *S. parvula*

Prior to this study, the *S. parvula* reference gene models (v2.2) were predicted based on *ab inito* methods as well as RNA-seq evidence based prediction derived from non-stressed conditions ^29^. To maximize the identification of transcripts that may be conditionally expressed under stress, we used 4-week-old *S. parvula* plants treated with multiple salts (NaCl, KCl, LiCl, and H_3_BO_3_) that are found at high levels in its native soils ^50^ for PacBio Iso-Seq sequencing. We obtained 500,265 error corrected circular consensus sequences (CCS) as our primary source of sequence reads to create an isoform specific reference transcriptome and to supplement the genome-based transcript annotation for *S. parvula.* We identified putative full-length transcripts based on 338,812 high quality CCS reads that contained 5’ and 3’ primers and polyA tails (Figure 1A). Following iterative clustering, error correction, and mapping to the *S. parvula* reference genome, we annotated 16,828 (corresponding to 11,348 genomic loci) structurally distinct putative protein coding transcripts expressed in *S. parvula* tissues exposed to multiple salts (see Methods for details). This added 7,732 new protein coding transcript models to the *S. parvula* reference genome to provide a total of 34,582 reference protein coding transcripts (Table 1).

**Figure 1.**
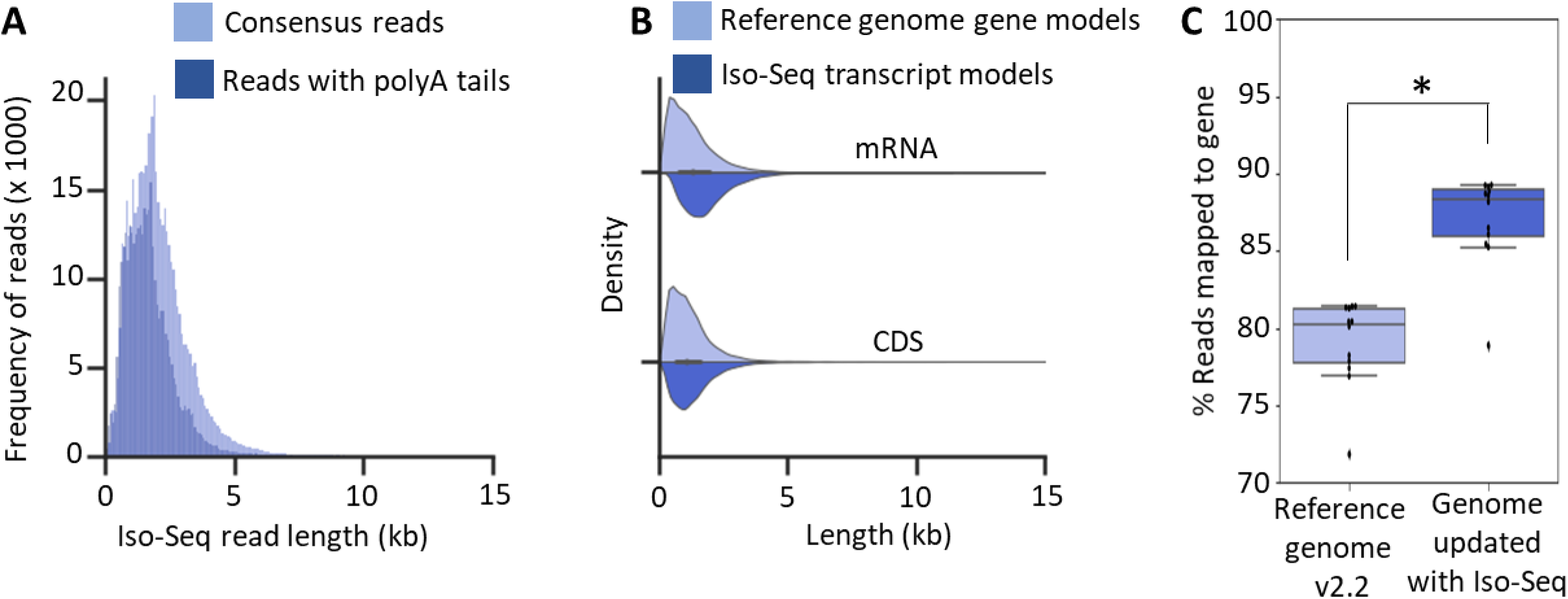
Improved *S. parvula* gene models using full length transcripts. **[A]** Majority of error corrected circular consensus reads (CCS) contain full length reads with polyA sequences and an average length of 1.7 kb. **[B]** Length distribution of Iso-Seq based transcript models and *S. parvula* Reference genome v2.2 gene models. **[C]** Percentage of mapped reads to the current genome and genome updated with Iso-Seq transcript models. Data = mean ± SD. Dots represent 4 independent RNA-Seq datasets used in this study as biological replicates (n = 12). Asterisk indicates significant difference (*p* ≤ 0.05), determined by one-sided t-test.

**Table 1.** Summary of *S. parvula* V2 genome updated with transcript models supported by Iso-Seq

| Category | Number of genes or transcripts |
| --- | --- |
| Genes supported by an Iso-Seq based transcript model | 11,348 |
| Total transcript models identified from Iso-Seq | 16,828 |
| Gene models identified with at least one new isoform | 7,028 |
| New genes identified from Iso-Seq | 301 |
| Gene models supplemented with UTR information using Iso-Seq | 11,348 |
| Genes with alternative splicing identified using Iso-Seq | 3,470 |
| Genes with alternative starts identified using Iso-Seq | 4,760 |
| Genes with alternative ends identified using Iso-Seq | 4,756 |
| Total number of protein coding transcripts annotated in the genome | 34,582 |

We were able to improve the *S. parvula* reference genome to include full length transcripts inclusive of 5’ and 3’ UTR regions with Iso-Seq reads. The average length of new Iso-Seq supported reference transcript models was greater than the corresponding length of transcript models in the *S. parvula* v2.2 reference genome annotation (Fig. 1B). The increase in transcript lengths was largely due to the identification of 5’ and 3’ UTR sequences that were previously missed in transcript model predictions in the reference genome. This refinement of reference transcript models generated UTR length distribution comparable to that of *A. thaliana* reference genome (Araport11) (Figure S1) and significantly increased the percentage of standard RNA-seq reads mapped to the reference transcriptome (Fig. 1C). This is expected to improve estimates of gene expression counts when using short-read RNA-seq data.

New genes previously not reported for *S. parvula* was added with Iso-Seq supported transcripts. The current reference *S. parvula* v2.2 genome includes 26,847 total protein coding primary gene models. The Iso-Seq supported transcripts mapped to 11,348 (42%) of those genes (Table 1). We additionally identified 301 novel gene models that were missed (i.e. sequence present in the genome but annotation absent) in the *S. parvula* reference genome. For example, the putative ortholog of the Arabidopsis *Magnesium/proton exchanger* (*MHX*, similar to *At2G47600)* in the *S. parvula* genome was annotated on chromosome 4 between *Sp4g29520* and *Sp4g29540,* using an Iso-seq based transcript model detected in this study (Figure S2). The novel transcript models further improved the reference genome annotation by adding multiple isoforms assigned to gene models, alternative transcription start and end sites for existing models, and UTR sequences (Table 1). The improved gene models, isoform specific expression, and Iso-Seq reads are available at Bioproject ID PRJNA63667.

We identified 5,911 alternatively spliced events, resulting in structurally different protein coding regions from the primary transcript models in the *S. parvula* genome from this study. These splice variants were categorized into intron retention, alternative 3’ acceptor, alternative 5’ donor, exon skipping, use of alternative first exon, use of alternative last exon, and use of mutually exclusive exons based on their frequency (Table 2). Intron retention was the most prevalent (55.2%) alternatively spliced event in *S. parvula.* We observed that two or more distinctly spliced isoforms could be co-expressed in either shoots or roots (Figure S3) when multiple isoforms were checked for their expression using RT-PCR for a select set of genes.

**Table 2.**
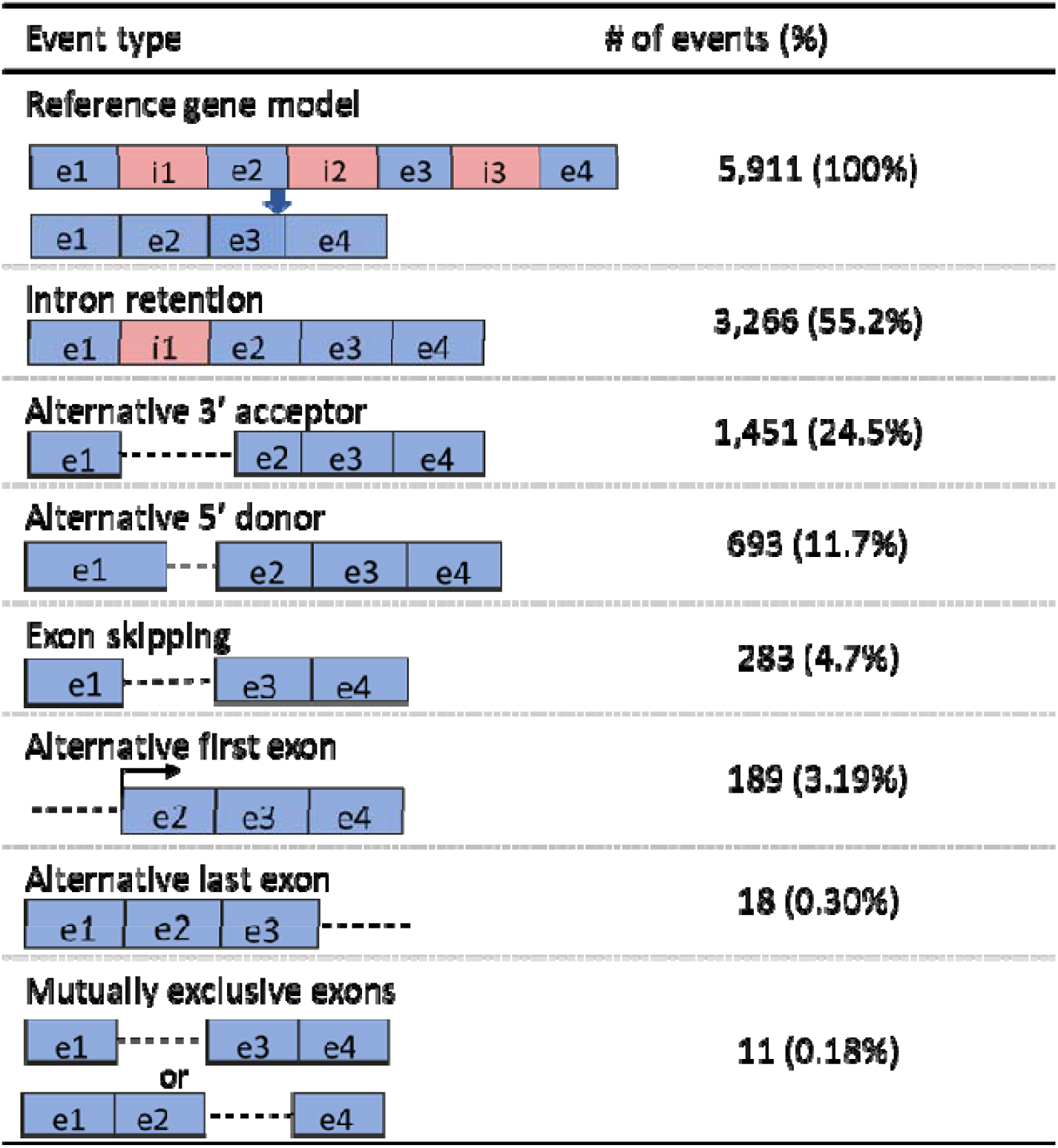
Major alternative splicing events identified using Iso-Seq reads in S. parvula. Blue boxes represent exons while red boxes or dash lines represent introns.

### Salt stress associated genes show a higher isoform diversity in *S. parvula* compared to *A. thaliana*

Alternative splicing can increase the repertoire of transcripts that are available to respond to abiotic stresses more efficiently and dynamically, independent of gene copy number variation ^51^. Therefore, we hypothesized that *S. parvula* would have a higher diversity of alternatively spliced isoforms for genes related to abiotic stress tolerance, specifically salinity tolerance, than in the less-tolerant species *A. thaliana*. To test this, we calculated the isoform ratio per ortholog pair in *S. parvula* and *A. thaliana* using the *S. parvula* reference isoforms identified in this study and *A. thaliana* reference isoforms obtained from AtRTD2 database ^16^. To avoid missing data or lack of expression of a certain gene in mature shoots or roots in one species being inferred as lack of isoform diversity in that species, we limited our comparison to genes expressed in our study that were represented by at least one transcript model in both species. We identified 10,859 *A. thaliana - S. parvula* ortholog pairs that had one or more isoforms per ortholog in each species (Figure 2A; Supplementary Table 2). Among them there were 6,874 ortholog pairs showing more isoforms in *A. thaliana* while only 1,201 pairs had a higher isoform number in *S. parvula* (Fig. 2A). Ortholog pairs annotated as “Response to stress” (GO:0006950) and “Transport” (GO:0006810) had a higher isoform diversity in *S. parvula,* while ortholog pairs annotated under “Nitrogen metabolism” (GO:0034641) had a higher isoform diversity in *A. thaliana* (Fig. 2B). As a control, we examined the distribution of isoforms in all ortholog pairs and found that these distributions were not significantly different between the two species (Fig. 2B).

**Figure 2.**
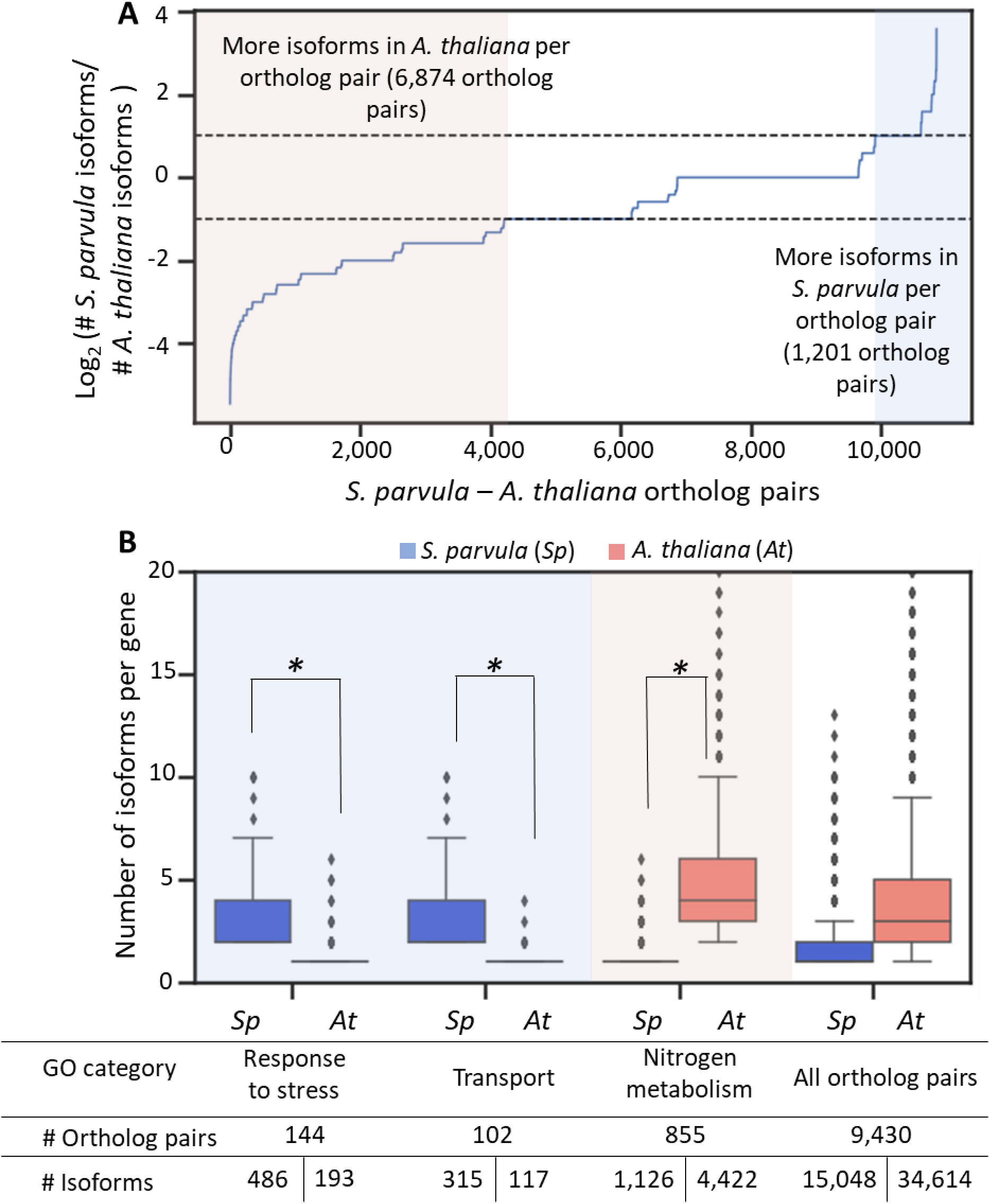
Genes with higher isoform diversity are associated with stress responses in *S. parvula* compared to *A. thaliana*. [A] Number of isoforms of *S. parvula* and *A. thaliana* per single-copy ortholog pairs given as a ratio. Blue and pink shaded areas indicate ortholog pairs where one species has more isoforms than the other. [B] Enriched functions associated with ortholog pairs that show at least 2-fold difference in isoform ratio between *S. parvula* and *A. thaliana.* Center line in the boxplots indicates median; box indicates interquartile range (IQR); whiskers show 1.5 × IQR. Asterisks indicate significant difference between isoform distributions of the two species, measured by Wilcoxon rank sum test at *p*-value cutoff ≤ 0.05.

The genes that had a higher isoform diversity in *S. parvula* included some of the most highly conserved and key stress responsive genes in plants including the Na^+^/H^+^ antiporter, *SOS1* known for its role in excluding Na^+^ from roots during salt stress ^52^ and *P5CS1* that codes for delta1-pyrroline-5-carboxylate synthase, the rate-limiting enzyme in proline biosynthesis known for its role in oxidative and osmotic stress responses ^53^. Notably, both *SOS1* and *P5CS1* are represented by single copy orthologs in *S. parvula* and *A. thaliana.* Five *SOS1* (out of 8 detected) and 8 *P5CS1* (out of 22 detected) isoforms for *S. parvula* were expressed at ≥ 0.5 TPM in both shoots and roots in control as well as salt treated conditions (Figure S4). The AtRTD2 database reported three *SOS1* and six *P5CS1* isoforms for *A. thaliana* ^16^.

### Isoform usage is less disordered in *S. parvula* compared to *A. thaliana* during salt stress

Diversity and conditional expression (i.e. specificity) of isoforms can be assessed using the Shannon entropy based information theory applied to transcriptomes ^54^. Stressed compared to growth optimal conditions are known to have higher transcriptome entropy and disorderdness with an increased number of alternative splice events when assessed using Shannon Entropy ^46^. We hypothesized that *S. parvula* transcriptomes will show a smaller entropy increase in its isoform usage when transitioning from control to salt stressed treatments compared to the salt-sensitive model *A. thaliana.* To test if isoform usage from control to stressed conditions went through a measurable entropy transition distinctive of the species, we used RNA-seq data from root and shoot samples to quantify the isoform abundance in *S. parvula* and *A. thaliana* and calculated the Shannon entropy (see Methods). We used *A. thaliana-S. parvula* ortholog pairs that were represented by at least two expressed isoforms with a normalized expression ≥ 0.5 TPM per ortholog within a species to avoid incomplete comparisons due to rare isoforms difficult to quantify in one species. This resulted in a total of 1,678 and 1,592 ortholog pairs expressed in roots and shoots. Roots had 3,832 and 5,239 isoforms for *S. parvula* and *A. thaliana* while shoots had 3658 and 4431 isoforms respectively. We found that both *S. parvula* and *A. thaliana* root transcript distributions increased mean entropy in response to salt stress (Figure 3A). This is aligned with the expectation that stress conditions create higher transcript diversity, lower specificity, and more disorderdness in transcript expression compared to a stress-neutral control condition ^55^. *A. thaliana* shoots showed a significant increase in entropy when transitioning from control to salt stressed conditions (Fig. 3A). The change in entropy for *S. parvula* was less in both roots and shoots suggesting a less disordered state of isoform usage compared to the relatively stress-sensitive *A. thaliana* when responding to stress conditions. We observed that the isoform usage in response to salt was highly species specific. The number of ortholog pairs that showed increased or decreased isoform usage as a shared response to salt stress in both species roots (156 expressed orthologs) and shoots (142 expressed orthologs) were much fewer than those orthologs (977 in roots and 937 shoots) that had a specific usage change in one species (Fig. 3B). Orthologs that showed high isoform usage specificity (i.e. maintained or lowered entropy) in response to salt stress in *S. parvula* roots compared to *A. thaliana* were enriched in functions largely associated with salt stress (Fig. 3C). In shoots, genes that maintained isoform usage specificity under salt stress in both species were enriched in salt stress associated functions (Fig. 3C).

**Figure 3.**
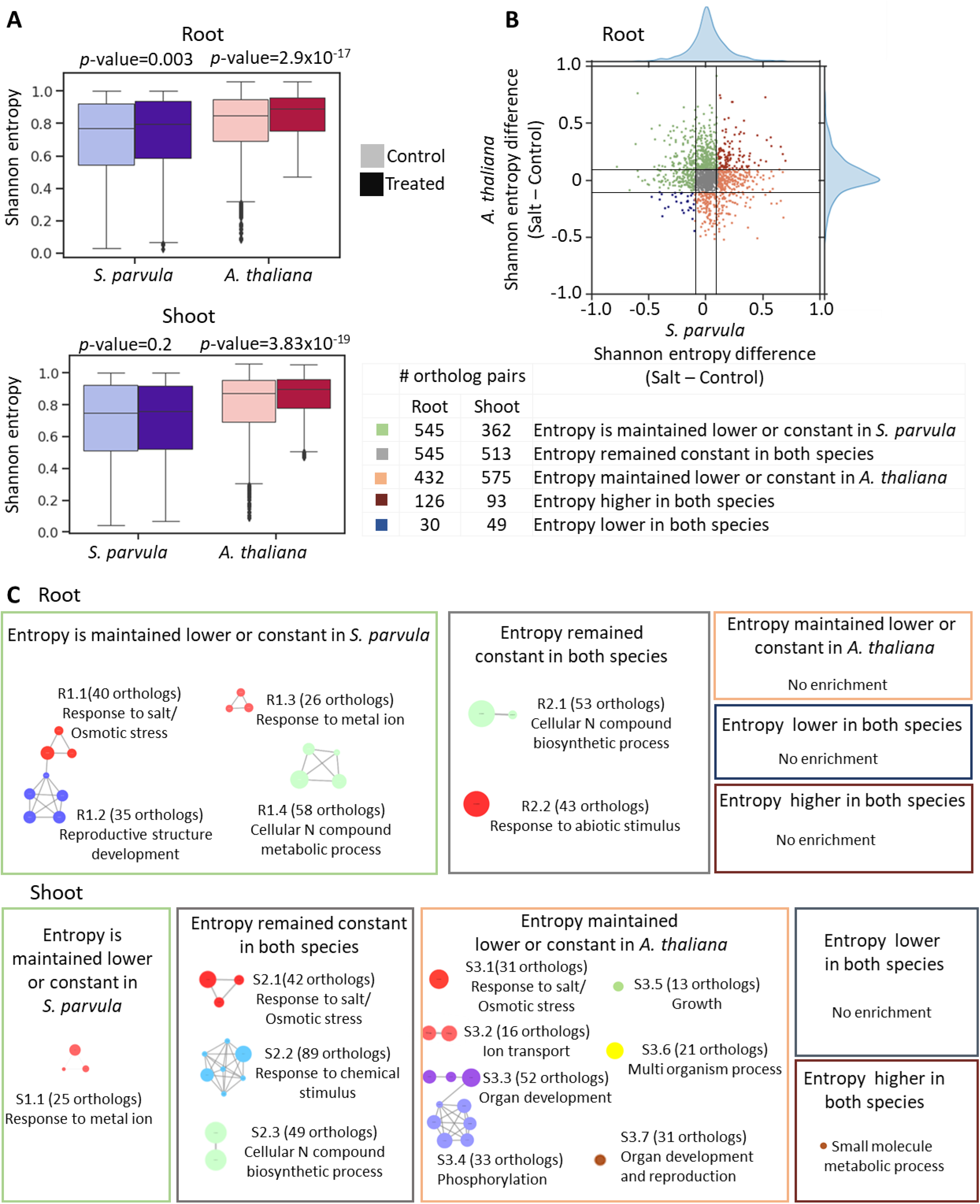
Isoform usage specificity between control and salt treated conditions. **[A]** Shannon entropy distribution of 1,678 ortholog pairs with at least two isoforms expressed per ortholog per species. Center line in the boxplots indicates median; box indicates interquartile range (IQR); whiskers show 1.5 × IQR. Each treatment was compared to control according to Student’s t test with *p*-values indicated above the relevant pairs. **[B]** Shannon entropy change between salt and control conditions in *S. parvula* and *A. thaliana* ortholog pairs in roots. Each dot represents an ortholog pair. Black lines indicate 0.5 entropy differences. Frequency distribution of data are shown on the marginal plot. **[C]** Functionally enriched processes represented by ortholog pairs in distinct categories of entropy shifts. A node in each cluster represents a gene ontology (GO) term; size of a node represents the number of genes included in that GO term; the clusters that represent similar functions share the same color and are given a representative cluster name and ID; and the edges between nodes show shared genes between functions. All clusters included in the network have adj *p*-values ≤0.05 with false discovery rate correction applied.

### Distinct regulation between different isoform usage and differential expression in response to salt stress

We next examined if the differently expressed genes in response to salt stress were also subjected to changes in their isoform usage under high salinity. Supplementary Table 3 lists all genes identified as differently spliced or differently expressed. Genes that were differently expressed as well as differently spliced in response to salt stress were rare in *S. parvula* and *A. thaliana* (≤ 3%) (Figure 4A). Moreover, the shared orthologs that were differently spliced in response to salt stress between species either in roots or shoots were also low (~3%) (Fig. 4B). Multiple genes differently expressed under salt stress in *A. thaliana* are found to be only differently spliced in response to salt in *S. parvula* (Supplementary Table 3). Figure S5 further highlights the high degree of species-specific regulation in differential isoform usage in response to stress. However, there is high convergence in the enriched functions represented by differently spliced isoforms in response to salt stress in both species (Fig. 4C).

**Figure 4.**
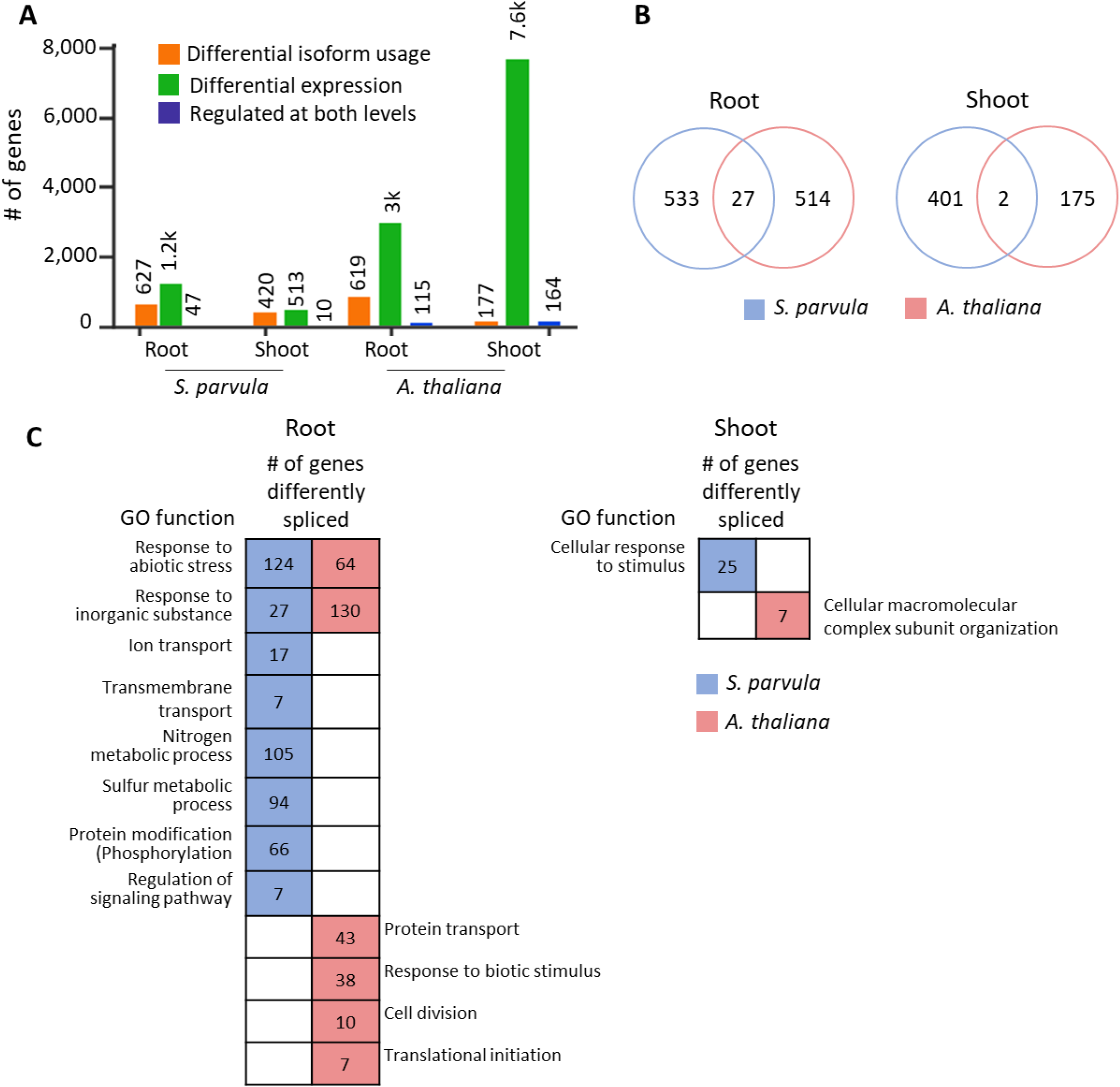
Genes differently spliced and differently expressed in response to salt stress. **[A]** Number of genes in *S. parvula* and *A. thaliana* that are differently regulated under salt stress. **[B]** Number of orthologs that show differential splicing in *S. parvula* and *A. thaliana* root and shoot in response to salt. [**C**] Functionally enriched processes represented by differently spliced genes in *S. parvula* and *A. thaliana*.

### Non-canonical splice sites are enriched in stress associated genes

Majority of splice sites in plants are marked by GU at the 5’ and AG at the 3’ sites in introns ^56^. Although less common, plant genes are spliced at alternative sites termed as non-canonical splice sites and alternative splicing at non-canonical sites are associated with abiotic stress responses ^56–58^. We investigated whether the expression of transcripts with non-canonical splice sites (Supplementary Table 4) increased under salt stress in *S. parvula* compared to *A. thaliana.* We found that *S. parvula* did not show any significant difference in mean expression strength between non-canonical and canonical transcripts in both roots and shoots while *A. thaliana* shoots showed an increased expression in transcripts that had non-canonical splice sites when treated with salt (Figure 5A).

**Figure 5.**
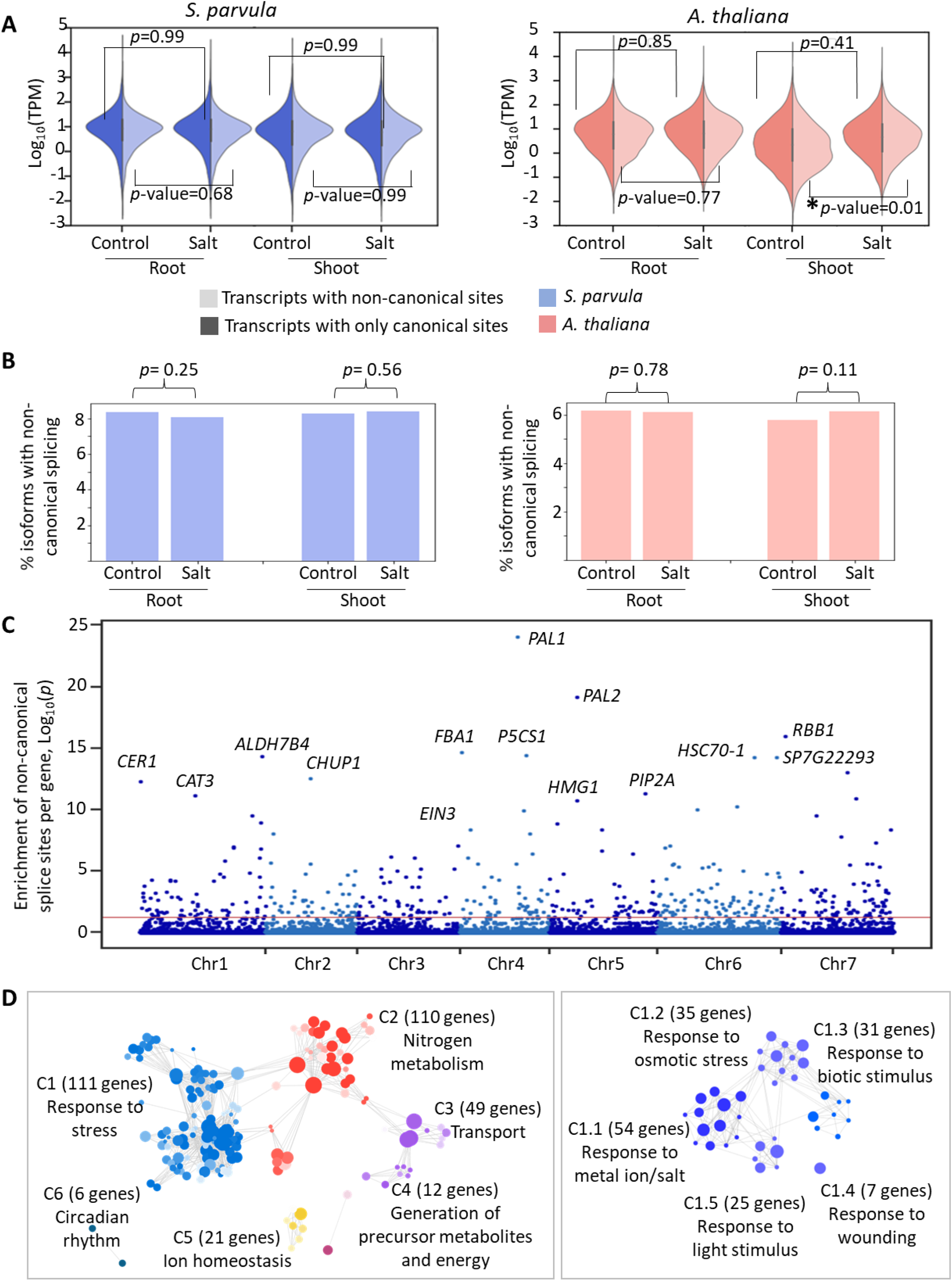
Use of non-canonical splice sites in transcripts expressed under stress. **[A]** Expression distribution of transcripts that contain only canonical splice sites and transcripts with at least one non-canonical splice site in roots and shoots of *S. parvula* (left panel) and *A. thaliana* (right panel). Asterisks indicate significant difference (*p* ≤ 0.05) of expression distributions between control and salt treated condition measured by two-sided t-test. **[B]** Number of expressed non-canonically spliced transcripts as a % out of total transcripts expressed in *S. parvula* and *A. thaliana* in response to salt. Significant differences between control and stress conditions were tested using Fisher’s exact test. **[C]** *S. parvula* genes that were enriched in non-canonical splice sites. The y axis shows the −log_10_p-value for a test of excess of non-canonical splice sites computed using a binomial test, where the probability of enrichment is calculated as the total non-canonical splice sites divided by the total number of splice sites per gene, ordered in the chromosomal order (x-axis) for the *S. parvula* genome. Genes with a high enrichment for non-canonical splicing are labeled. Red line indicates the −log_10_p corresponding to adjusted *p*-value of 0.05. **[D]** Functional processes enriched in genes detected to be non-canonically spliced in [C]. A node in each cluster represents a gene ontology (GO) term; size of a node represents the number of genes included in that GO term; the clusters that represent similar functions share the same color and are given a representative cluster name and ID; and the edges between nodes show the connectivity of genes between functions. All clusters included in the network have adj *p*-values ≤0.05 with false discovery rate correction applied. More significant values are represented by darker node colors. The right panel shows the sub-clustered functions represented by the largest cluster C1 in the left panel.

Previous studies have reported increases in non-canonical splicing in plants under abiotic stresses ^59,60^. Therefore, we examined if usage of transcripts with non-canonical splice sites significantly increased under salt stress compared to control conditions in *S. parvula* differently from *A. thaliana.* Similar to previous reports, non-canonically spliced transcripts are less frequent than canonically spliced transcripts regardless of the condition tested (≤ 10%; Fig. 5B). However, our analysis does not find a significant increase in non-canonically spliced isoforms from control to salt treated conditions in either species (Fig. 5B).

Next, we tested if genes with non-canonical splice sites were enriched for stress associated functions in *S. parvula.* We found 424 genes out of 25,145 multi-exon coding genes to be enriched in non-canonical splice sites in the *S. parvula* genome (Fig. 5C). Some of these are notable genes associated with stress regulatory pathways (for example, *PAL1, PAL2, P5CS1* and *HSC70-1*) (Fig. 5C)*. S. parvula* genes enriched for non-canonical splice sites were indeed primarily enriched in stress response pathways (Fig. 5D). Further sub-clustering of the functional group annotated under “stress responses” (cluster C1 of Fig. 5D) showed that genes in salt/metal ion and osmotic stress were specifically contributing to this cluster.

### Predicted isoforms for the *S. parvula* genome is enriched for stress responsive genes

It is likely that we may have missed to detect stress responsive isoforms expressed in *S. parvula* in this study because exhaustive searches for conditionally expressed isoforms are impractical for emerging model organisms. Therefore, we sought to employ a machine learning approach to predict alternative splicing sites in the *S. parvula* genome as an alternative. We applied the deep neural network, SpliceAI which is expected to yield high confidence predictions among recent tools developed to predict splice events using genomic sequences ^48,61,62^. We used the known splice site information from *A. thaliana* chromosomes 1-4 to train the SpliceAI network and received an average precision of 0.92 when tested with *A. thaliana* chromosome 5 (Figure 6A). We then predicted splice sites from 26,847 *S. parvula* pre-mRNA sequences and obtained 214,901 splice site predictions including 114,284 novel splice sites (Fig. 6B). Twenty-six percent of splice sites previously observed were also predicted using SpliceAI and we found 7,302 genes with at least one newly, predicted isoform. Prediction probability scores were highest for splice sites within the gene compared to those in the first and the last introns (Figure S6).

**Figure 6.**
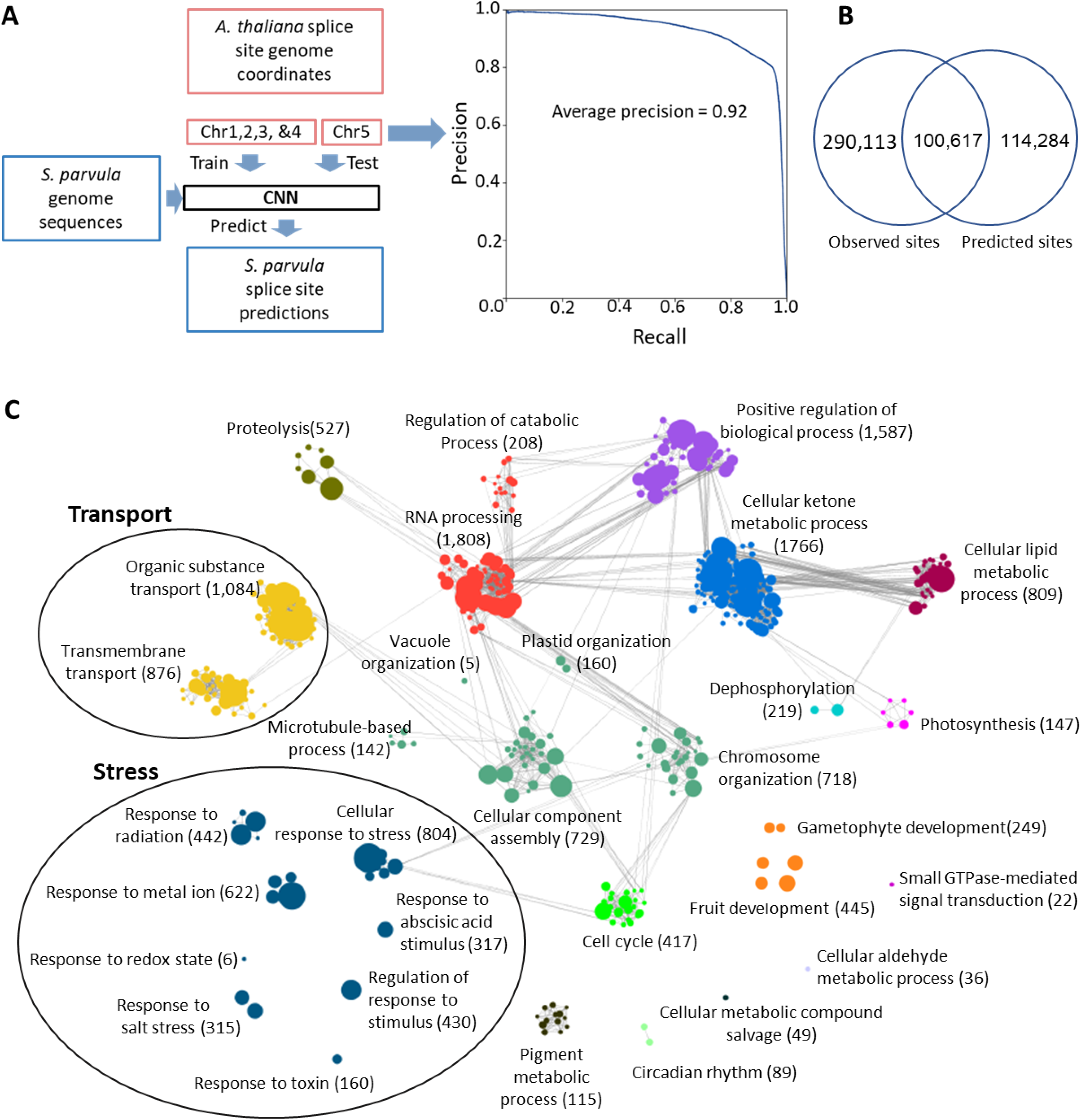
Genome wide prediction of splice sites for *S. parvula* using a deep neural network. **[A]** Training and testing with *A. thaliana* and application of the SpliceAi model to *S. parvula.* **[B]** Overlap between the observed and predicted splice sites for *S. parvula* protein coding gene models. A probability score ≥ 0.6 was used for the predicted splice sites. **[C]** Functional processes enriched in genes observed and predicted to have more than one isoform in the *S. parvula* genome. A node in each cluster represents a gene ontology (GO) term; size of a node represents the number of genes included in that GO term; the clusters that represent similar functions share the same color and are given a representative cluster name; and the edges between nodes show the connectivity of genes between functions. All clusters included in the network have adj *p*-values ≤0.05 with false discovery rate correction applied.

With the current analysis, we have identified 16,061 potential protein coding isoforms (observed or predicted) for 9,033 genes in the *S. parvula* genome (Supplementary Table 5). Interestingly, stress and transport associated functions are enriched among those genes that are observed or predicted to have more than one isoform (Fig. 6C). Stress and transport related functions deduced from GO annotations account for 35% of genes that are alternatively spliced in the *S. parvula* genome.

## Discussion

### Alternative isoforms of stress related genes from an extremophyte model as a resource in environmental stress adaptations

Alternative splicing allows genes to acquire new functions independent from gene duplications and promoter evolution. Previous studies have shown that duplicated genes are enriched in stress associated functions in *S. parvula* and other extremophytes facilitating their stress adapted lifestyles more than in stress-sensitive sister species ^29,63,64^. However, extremophyte gene diversity represented by alternatively spliced isoforms is underexplored ^65^. Certain genes are regulated only at the alternative splicing level with no change at the gene expression level that have led to the increasing recognition of the importance of isoform specific reference transcript datasets in gene expression studies^66^. In this study we examined the possibility of diversifying gene functions through alternative splicing and specially focused on isoforms differently used during salt stress in one of the leading model extremophytes ^26^.

*Schrenkiella parvula* and *A. thaliana* genomes have similar gene numbers (~27,000) and similar genome sizes (~120 MB) ^29^. A recent study that explored alternative splicing in *A. thaliana* using full length transcript sequencing based on Iso-Seq reports the discovery of isoforms in similar proportions to our study with intron retention being the most common alternative splicing event ^67^ (Table 2). This suggests that *S. parvula* is not an exception in highly increased or decreased transcript diversity through alternative splicing although the recorded number of isoforms for the model plant through aggregate studies using multiple tissues, developmental stages, and treatments are much higher (Zhang et al., 2017). Given the genomic similarities between *S. parvula* and *A. thaliana,* their transcriptome adjustments with differential splicing in response to salt stress were remarkably distinct from one another when an identical salt treatment was given to mature plants (same age and tissues tested in both species) (Figs. 4 and S5).

In support of our hypothesis that extremophytes would diversify their response to stress via alternative splicing in selected gene groups, we observed that *S. parvula* orthologs had a higher number of isoforms compared to *A. thaliana* in genes associated with stress and transport functions (Fig. 2). Stress and transport functions were also enriched among duplicated genes in *S. parvula* compared to *A. thaliana ^68^.* We found that differently expressed genes and genes that showed differential isoform usage were largely mutually exclusive within species as well as in one-to-one ortholog pairs between *S. parvula* and *A. thaliana* (Figs. 4 and S5). Further, when we combine both observed and predicted splice sites in the *S. parvula* genome, the potential protein coding isoform pool is enriched in functions associated with stress tolerance (Fig. 6). These observations together indicate that genes expressed in response to stress are highly diversified and non-overlapping in their mode of function, but converge on common functions associated with stress tolerance in *S. parvula.* Therefore, our study provides a novel resource for assessing functional significance of stress tolerance genes in the extremophyte model. It allows selection of target genes that could be tested at the isoform level when expression modulation via promoter modifications or single gene-knockouts of essential genes do not offer optimal methods to test novel gene functions contributing to stress tolerance.

### Isoform usage and the specificity of their expression in response to salt

Our current study in agreement with a previous study on *A. thaliana* have shown that most differently spliced genes were not differently expressed in response to salt stress representing an independent layer of gene regulation in response to stress ^21^. Compared to animals, plants tend to use alternative splicing biased to environmental stress responses more than for tissue-specific responses ^69^. Multiple studies have reported specific associations of alternative splicing and environmental stress in plants ^22,57,59,70,71^. However, fewer studies have examined the presence of non-specific alternative splicing leading to increased number of differently spliced isoforms under abiotic stress ^12,72^. Additionally, components of the spliceosome are differently expressed leading to differential splicing of target genes in *A. thaliana* during stress conditions ^60^. In animals, stressed conditions are reported to have increased amount of alternatively spliced isoforms with high non-specific expression, thus creating a higher level of disorderdness in isoform expression ^46^ which can be quantified using Shannon entropy ^73,74^. We predicted that plants will show a similar trend in increased disorderdness in isoform expression at the transcriptome level during stress conditions. Furthermore, we expected to see a smaller change in entropy in the extremophyte when transitioning to a salt treated condition compared to the stress-sensitive species. Indeed, this prediction was supported by the shoot transcriptomic response we observed for *S. parvula* and *A. thaliana* (Fig. 3). Notably, the genes that shifted to lower entropy values representing shifts to specific isoform in their expression specificity under stress were enriched for stress associated functions in both roots and shoots in *S. parvula* (Fig. 3). Our study cannot test if the tendency to increase transcriptome disorderdness via less specifically expressed isoforms per gene is indicative of aberrant splicing under stress. Yet, the comparison between *S. parvula* and *A. thaliana* suggests that the extremophyte is more prepared to respond to salt stress by specific isoforms mostly expressed for stress associated genes.

In conclusion, this study provides a novel resource for a leading extremophyte model and expands our knowledge on the ability to respond to stress via differential isoform usage independently from differential gene expression. Stress associated functions were enriched among genes observed or predicted to have multiple isoforms in *S. parvula;* one-to-one orthologs where *S. parvula* has a higher number of isoforms than *A. thaliana;* genes that showed differential isoform usage in response to stress in *S. parvula; S. parvula* genes that were enriched in non-canonical splice sites; and *S. parvula* genes that maintained or lowered their disorderdness by expression of specific isoforms under stress. These findings contribute to how we understand stress tolerance evolved in an extremophyte. Differential isoform usage offers a complementary path to increase the coding potential of the *S. parvula* genome that cannot be fully explained by gene duplication or promoter evolution alone. Future studies on other extremophytes exploring isoform diversity will facilitate the identification of convergent traits in isoform usage evolved in stress-adapted plants. Such a resource will be influential in deducing diverse stress responsive networks and identifying transferable stress responsive genes into crops.

## Acknowledgements

We thank Drs. Guannan Wang, Pramod Pantha, Dong-Ha Oh, Aaron Smith, John Larkin and graduate student Richard S Garcia for providing feedback on the manuscript and facilitating helpful discussions. This work was supported by the US National Science Foundation awards MCB-1616827 and IOS-EDGE-1923589, and US Department of Energy BER-DE-SC0020358 awarded to MD. CW and KT were additionally supported by an Economic Development Assistantship award from Louisiana State University. We acknowledge LSU High Performance Computing services for providing computational resources for this study.

## Author Contributions

CW and KT conducted wet lab experiments. CW performed bioinformatics analyses. MD developed the experimental design and supervised the overall project. CW, KT, and MD interpreted results and wrote the article.

## Figure legends

**Figure S1. UTR length distribution of transcript models in *S. parvula* and *A. thaliana.* [A]** 5’ UTR and **[B]** 3’ UTR length distributions. The distributions were obtained from 16,828 *S. parvula* and 41,064 *A. thaliana* transcript models

**Figure S2. Novel gene *SP4G29525 (SpMHX1)* annotated using Iso-Seq transcript models.** Two transcript models were detected with Iso-Seq full length reads for both *MHX* and *PHR2* gene models.

**Figure S3. Independent detection of selected isoforms first identified with Iso-Seq in *S. parvula* transcriptome. [A]** Transcript models of selected genes with isoform IDs. Coding region and UTR regions are indicated in dark and light shades. Locations of primer binding sites are shown by arrows. Exons are numbered. Introns are given as connecting lines between exons. Identical exon structures past the 2^nd^ exon between isoforms are represented by dashed lines. **[B]** Predicted protein coding regions and functional domains of the corresponding transcripts. Functional domains for each transcript are marked as colored blocks. **[C]** Gel electrophoresis images of amplified transcripts obtained from RT-PCR using primers indicated in A. Arrow heads indicate the expected size of the amplified product.

**Figure S4. *SOS1* and *P5CS1* isoform diversity in *S. parvula.* [A]***SpSOS1* isoforms expressed above 0.5 TPM in all conditions. *SpSOS1 (v2)* serves as the primary gene model annotated in the current genome annotation. **[B]** *SpP5CS1* isoforms expressed above 0.5 TPM in all conditions. *SpP5CS1 (v2)* serves as the primary gene model annotated in the current genome annotation. Data = mean ± SD (n = 3).

**Figure S5. Differential regulation of orthologs in *S. parvula* and *A. thaliana* in response to salt stress. [A]** Root and **[B]** shoot. UpSet plot numbers represent number of orthologs. DS - Differently spliced; DE – Differently expressed; Sp - *S. parvula*; At - *A. thaliana*.

**Figure S6. Probability of splice sites identified using SpliceAi from 5’ to 3’ for *S. parvula* gene models**.

Dashed line indicates the probability thresholds used to predict a splice site.

**Supplementary Table 1.** Primers used for RT-PCR.

**Supplementary Table 2.** Ortholog pairs between *S. parvula* and *A. thaliana* isoforms, annotations, and isoform ratio.

**Supplementary Table 3.** Differently spliced and differently expressed genes in *S. parvula* and *A. thaliana.*

**Supplementary Table 4.** Enrichment for non-canonical over canonical splice in *S. parvula.*

**Supplementary Table 5.** Predicted and observed splice sites in the *S. parvula* genome. Link: https://github.com/wchathura/Iso-Seq_Dataset/blob/main/Supplemntry_table_5.txt

