## Supplementary Figures for "Alternative splicing preferentially increases transcript diversity associated with stress responses in the extremophyte *Schrenkiella parvula*"

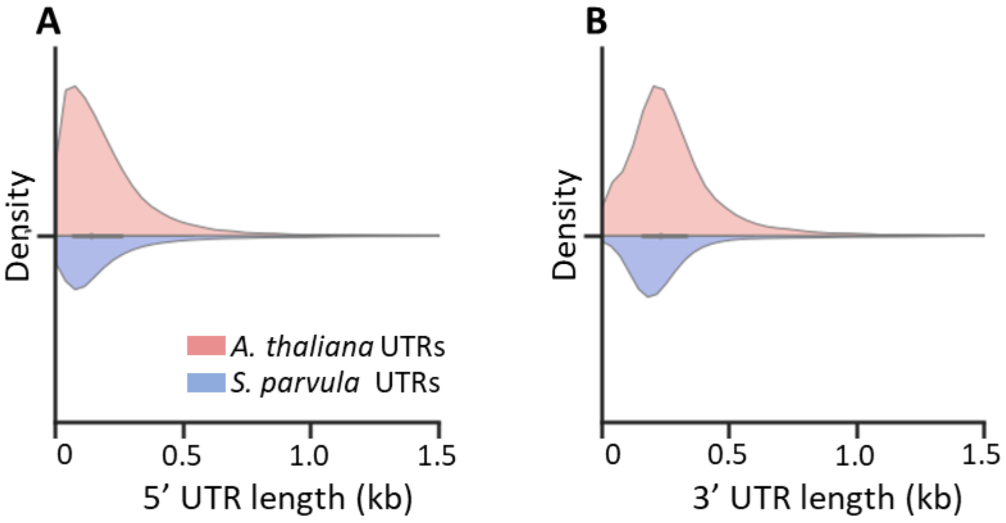


**Figure S1. UTR length distribution of transcript models in *S. parvula* and *A. thaliana*. [A]** 5′ UTR and **[B]** 3′ UTR length distributions. The distributions were obtained from 16,828 *S. parvula* and 41,064 *A. thaliana* transcript models.

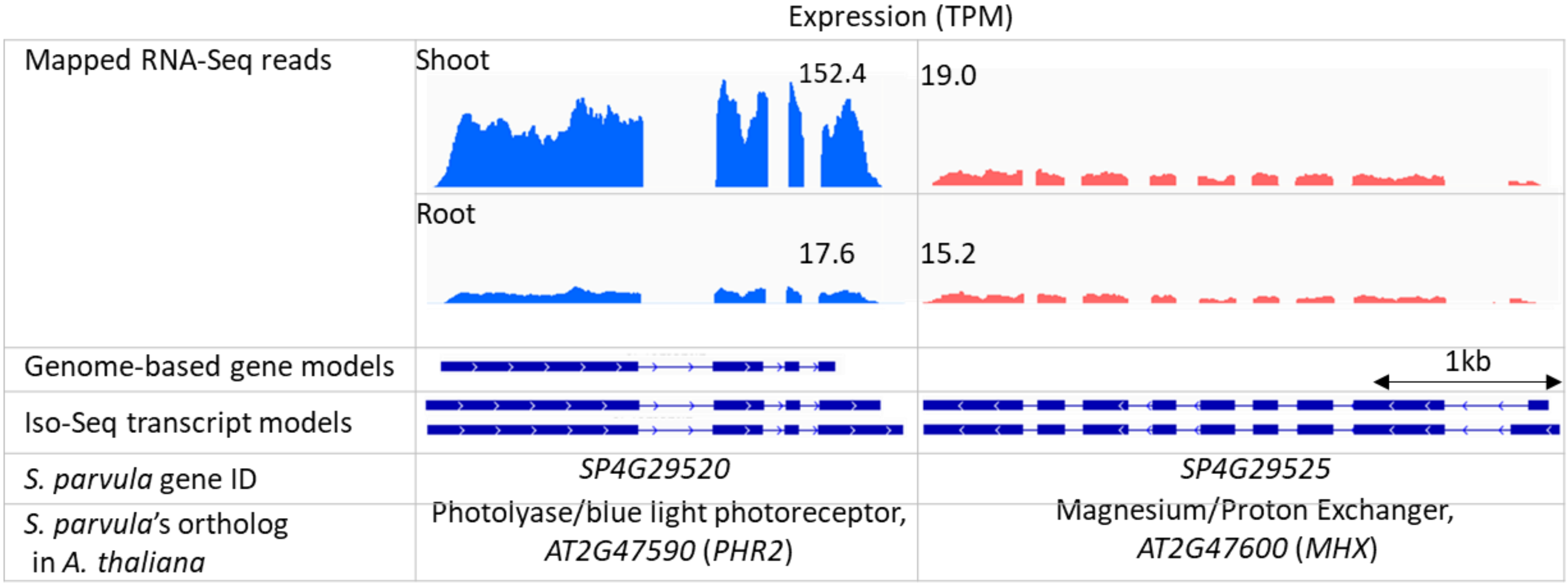


**Figure S2. Novel gene *SP4G29525 SpMHX1* annotated using Iso-Seq transcript models.** Two transcript models were detected with Iso-Seq full length reads for both *MHX* and *PHR2* gene models.

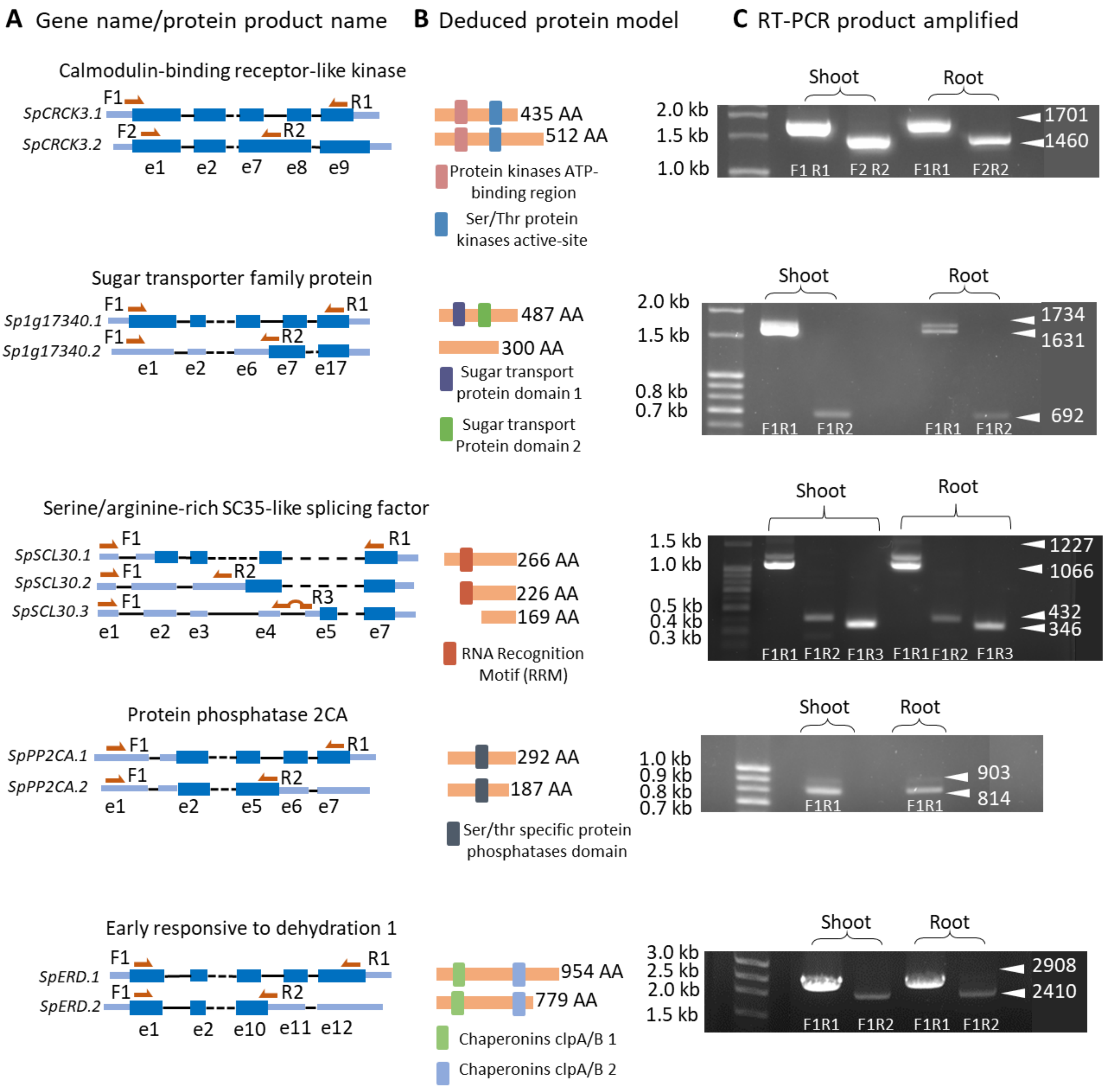


**Figure S3: Independent detection of selected isoforms first identified with Iso-Seq in *S. parvula* transcriptome. [A]** Transcript models of selected genes with isoform IDs. Coding region and UTR regions are indicated in dark and light shades. Locations of primer binding sites are shown by arrows. Exons are numbered. Introns are given as connecting lines between exons. Identical exon structures past the 2nd exon between isoforms are represented by dashed lines. **[B]** Predicted protein coding regions and functional domains of the corresponding transcripts. Functional domains for each transcript are marked as colored blocks. **[C]** Gel electrophoresis images of amplified transcripts obtained from RT-PCR using primers indicated in A. Arrow heads indicate the expected size of the amplified product.

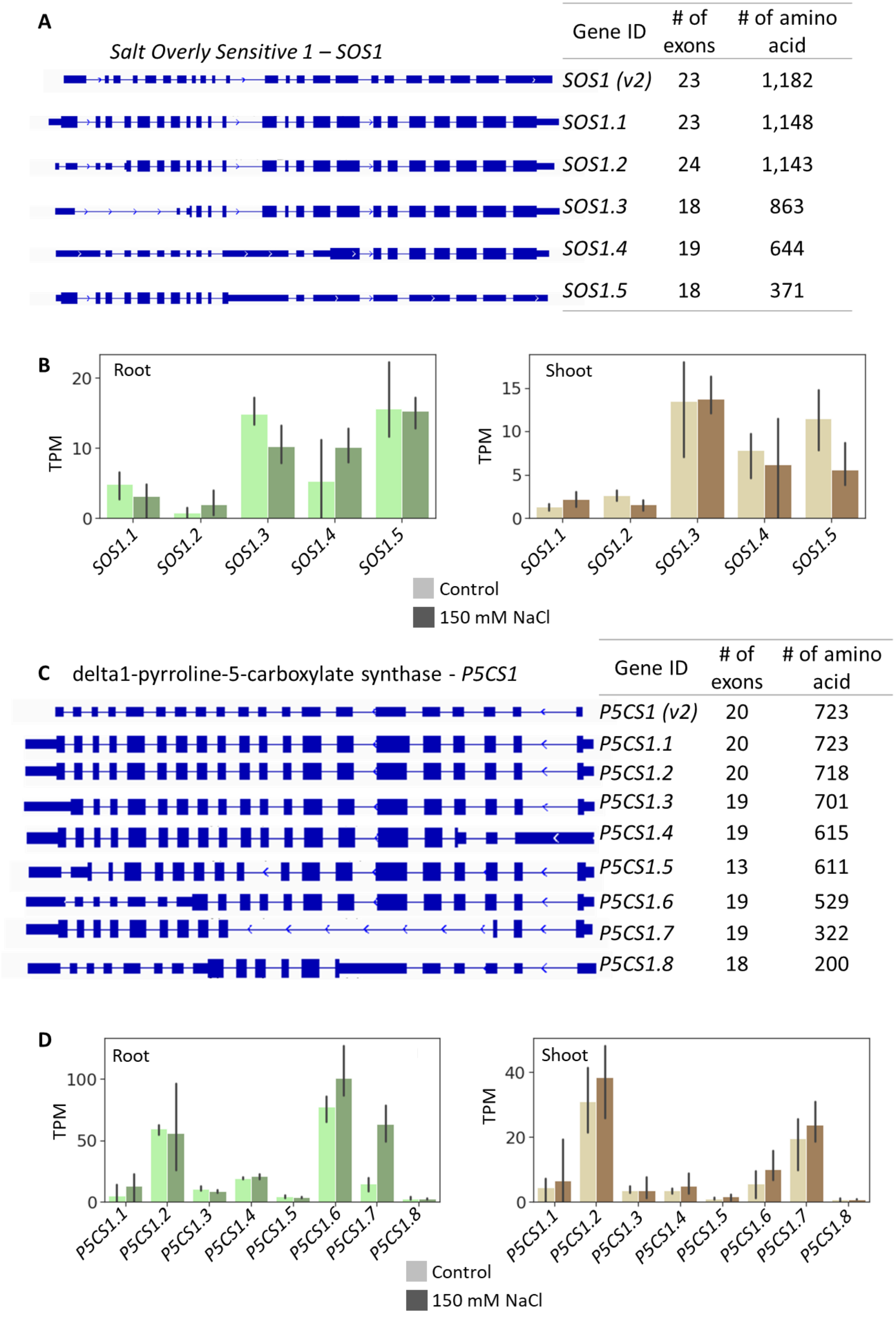


**Figure S4. *SOS1* and *P5CS1* isoform diversity in *S. parvula*. [A]** *SpSOS1* isoforms expressed above 0.5 TPM in all conditions. *SpSOS1 (v2)* serves as the primary gene model annotated in the current genome annotation. **[B]** *SpP5CS1* isoforms expressed above 0.5 TPM in all conditions. *SpP5CS1 (v2)* serves as the primary gene model annotated in the current genome annotation. Data = mean ± SD (n = 3).

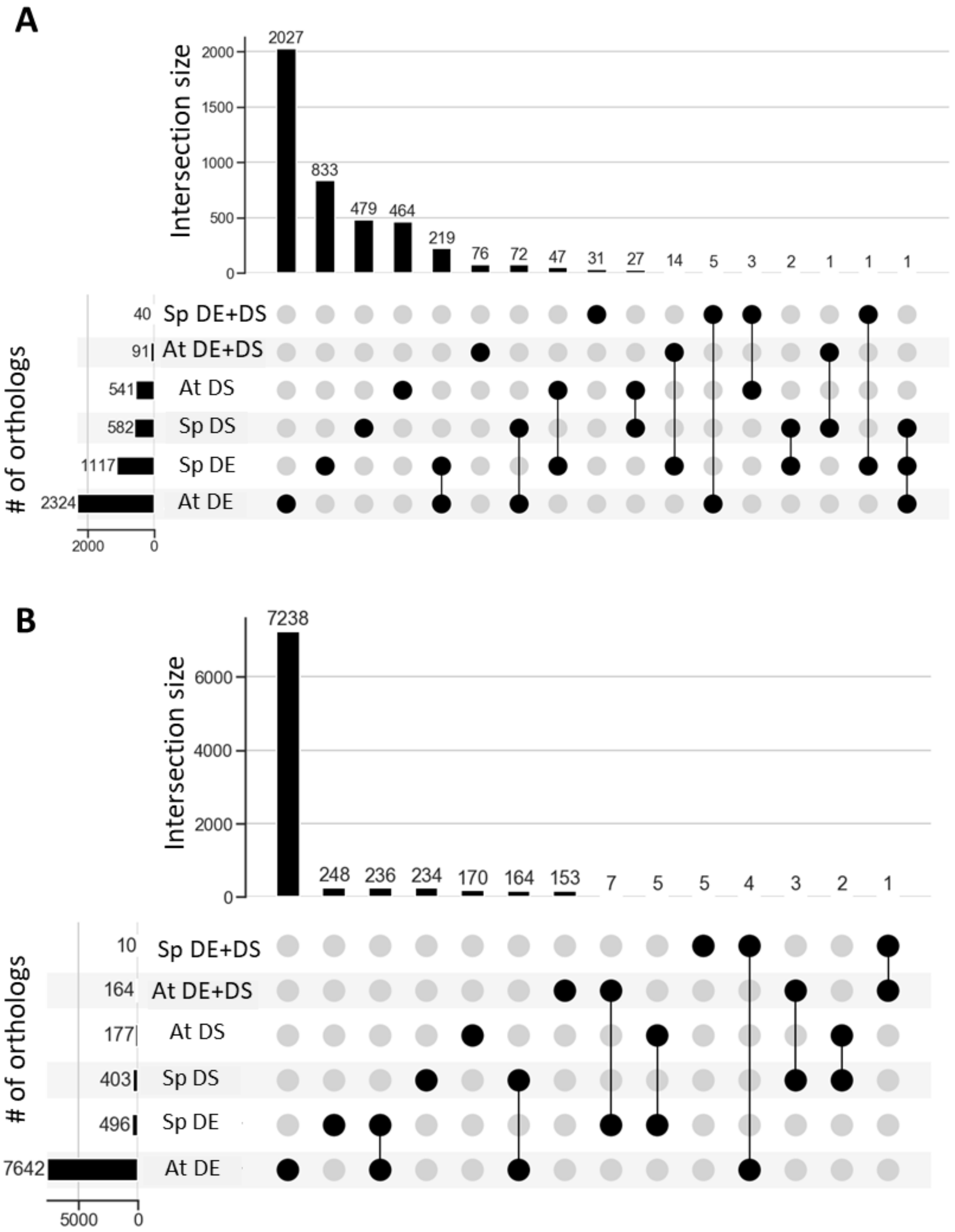


**Figure S5: Differential regulation of orthologs in *S. parvula* and *A. thaliana* in response to salt stress. [A]** Root and **[B]** shoot. UpSet plot numbers represent number of . DS - Differently spliced; DE – Differently expressed; Sp - *S. parvula;* At - *A. thaliana*.

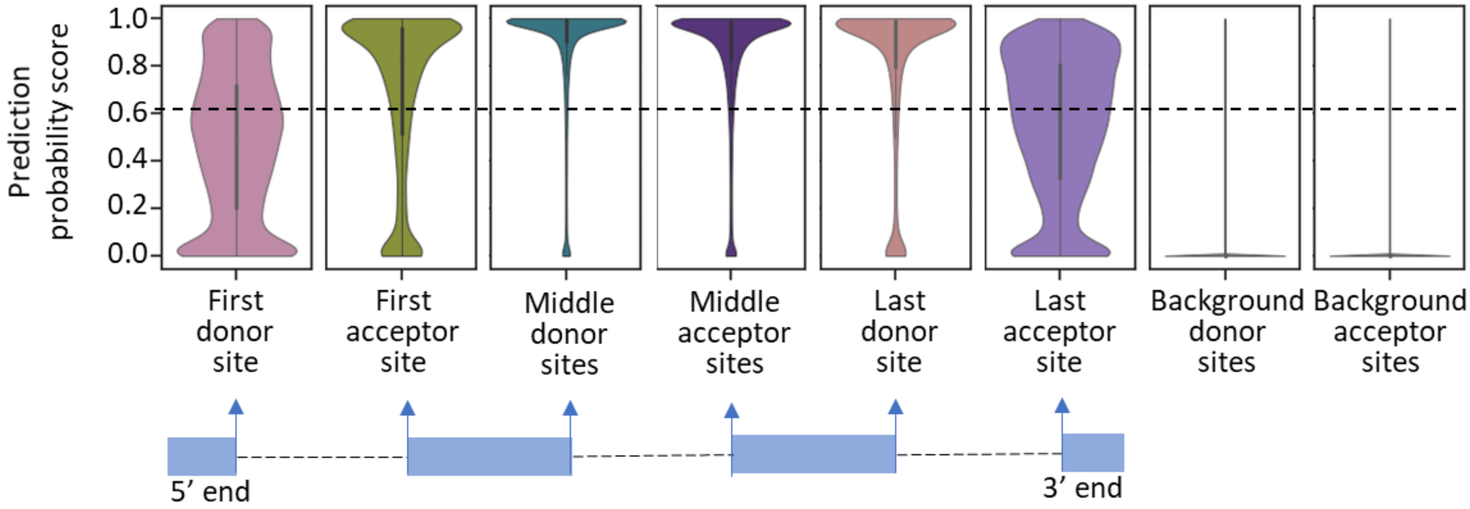


**Figure S6. Probability of splice sites identified using SpliceAi from 5' to 3' for *S. parvula* gene models**. Dashed line indicates the probability thresholds used to predict a splice site.
